# Environmental (e)DNA detection of the invasive pink salmon *Oncorhynchus gorbuscha* during the 2017 Norwegian invasion

**DOI:** 10.1101/651554

**Authors:** Laura M. Gargan, Frode Fossøy, Tor A. Mo, Jeanette E. L. Carlsson, Bernard Ball, Jens Carlsson

## Abstract

The pink salmon *Oncorhynchus gorbuscha* was introduced from its native range in the Pacific to Northwest Russia several times since the 1950’s. While this species has been regularly observed in rivers in Northern Norway since that time, there has been an upsurge in the numbers of odd-year *O. gorbuscha* individuals observed in rivers in southern Norway in recent years, and particularly in 2017. Although the wide-scale effects of this species presence are currently uncertain, there are concerns regarding potential competition between *O. gorbuscha* and native species – most notably the Atlantic salmon *Salmo salar.* Environmental (e)DNA is becoming a widely used tool to monitor rare and invasive species in aquatic environments. In the present pilot study, primers and a probe were developed to detect *O. gorbuscha* from eDNA samples taken from a Norwegian river system where the species was observed. Water samples were taken at both upstream and downstream locations of the Lysakerelva river during Autumn 2017 (to coincide with spawning) and during late Spring 2018. Autumn samples were positive for *O. gorbuscha* at both sampling locations, whereas Spring samples showed positive detection of this species in the upstream region of the river, when smolt should have left, or be in the process of leaving the river. These findings reveal that eDNA-based methods can be used detect the presence of *O. gorbuscha* during their spawning season. This suggests that odd-year populations have the potential to become established in the studied river system. We recommend that eDNA sampling is repeated to determine whether individuals of this odd-year population have survived at sea and return to spawn. Our assay specificity tests indicate that the tools developed in the present study can be used for detection of *O. gorbuscha* in both Norwegian and other European river systems where presence/absence data is required. We also suggest some modifications to our methodology that may improve upon the detection capabilities of *O. gorbuscha* using eDNA.

## INTRODUCTION

Pink salmon *Oncorhynchus gorbuscha* (Walbaum, 1792) are native to the Pacific Ocean, where they typically spawn in the freshwater ecosystems of bordering countries between the latitudes of 40o and 70o. This species follows a strict two-year life cycle (Figure 1). Adult *O. gorbuscha* migrate from the open sea and up-river to spawn in the autumn, after which all spawning individuals die. The juveniles emerge the following spring, ready to migrate down-river and out into the open sea to mature for one winter. They return as adults to freshwater in the next autumn to spawn, thus completing the life cycle (Heard 1991).

**Figure 1:**
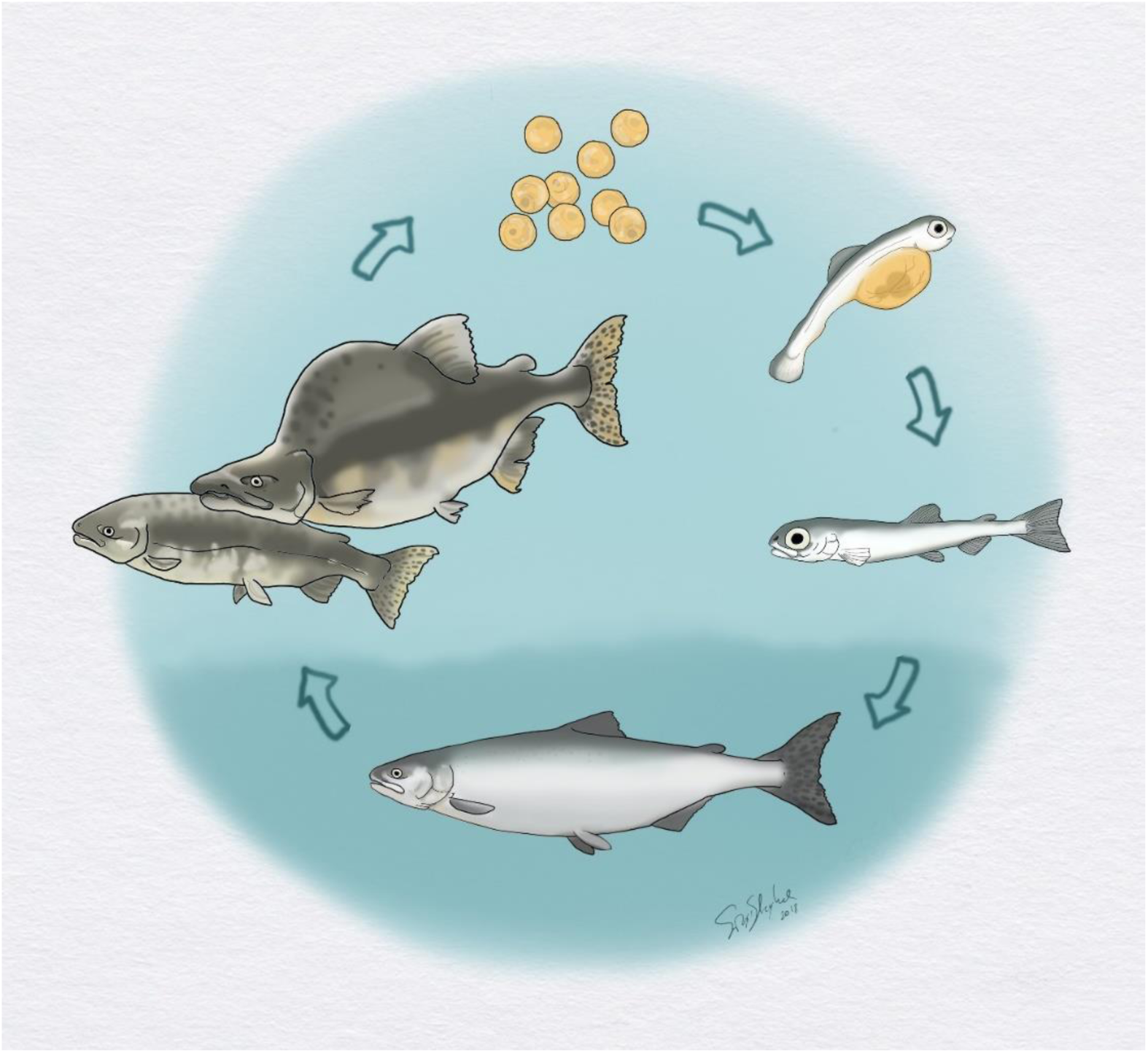
Life cycle of the pink salmon *Oncorhynchus gorbuscha, ill.: Sigrid Skoglund, NINA.*

*O. gorbuscha* was originally introduced from their native Pacific range to North Western Russia several times since the 1950’s, when fry were stocked in several rivers that drain into the White Sea and the Barents Sea (Bakshtansky 1980). While *O. gorbuscha* has been regularly found in rivers in Northern Norway since 1960 (Berg 1961), there has been an upsurge in the observations of odd-year *O. gorbuscha* in Norwegian rivers in recent years (Mo *et al.*, 2018), as well as in rivers in the UK and Ireland (Armstrong *et al.*, 2018, Whelan 2017, Millane *et al.*, 2019). However, self-reproducing populations have not yet appeared to become established, most likely because they have not adapted time of spawning to match local conditions outside of their native range (Mo *et al.*, 2018).

Established populations of *O. gorbuscha* in rivers in North Western Russia are dominated by odd-year individuals (Gordeeva & Salmenkova, 2011). Thus far, odd-year individuals have been noticeably more abundant throughout their invasive range, with a particularly large number of spawning *O. gorbuscha* recorded in 2017 (Armstrong *et al.*, 2018, Mo *et al.*, 2018). Therefore, it is these odd-year stocks that are most likely to become established as populations and the next spawning season (which should take place during Autumn of 2019) will be an important time for research scientists to determine the extent of this invasion and its potential ecological effects.

Presently, the spawning time of *O. gorbuscha* in Norway does not appear to overlap with native salmonids (e.g. Atlantic salmon *Salmo salar* and brown trout *S. trutta*). However, *O. gorbuscha* and *S. salar* have similar preferences for spawning habitats and so there is a risk of competition for optimal spawning sites (Sandlund *et al.*, 2018). While *O. gorbuscha* juveniles emerge ready to migrate to sea, observations in Norwegian rivers suggest that they spend some time feeding in freshwater (from weeks to months) and during this time there may be interactions between juveniles of native salmonids (Sandlund *et al.*, 2018). However, it is also possible that the eggs and fry of *O. gorbuscha* can provide a source of food for other native salmonid species (Rasputina *et al.*, 2016). In order to fully assess the impacts of the presence of *O. gorbuscha* in Norwegian (and other European) rivers, it is first necessary to determine the spatial, as well as the temporal (e.g. time of spawning and migration) distribution of this species.

Environmental DNA (eDNA) is a genetic survey method that relies on the detection of taxa from extracellular and intracellular material that is deposited into the environment. Subsequently, this material can be isolated from the environmental sample (such as water, air or soil; Taberlet *et al.*, 2012) and interrogated using genetic markers for multi-species (Thomsen *et al.*, 2012, Hänfling *et al.*, 2016) or targeted species (Ficetola *et al.*, 2008, Jerde *et al.*, 2011, Gustavson *et al.*, 2015) detection.

While many studies have used quantitative PCR (qPCR) for targeted detection of species in an eDNA sample (Thomsen *et al.*, 2012, Wilcox *et al.*, 2013, Atkinson *et al.*, 2018), more recently the approach of using digital droplet PCR (ddPCR) has been adopted by some researchers (Doi *et al.*, 2015, Nathan *et al.*, 2015, Evans *et al.*, 2017, Baker *et al.*, 2018), with either similar (Nathan *et al.*, 2015) or increased (Doi *et al.*, 2015, Hunter *et al.*, 2017) sensitivity reported for ddPCR platforms compared to qPCR for the analysis of eDNA. ddPCR is a relatively recent technological advancement, allowing for accurate estimation of low copy DNA number (Hindson *et al.*, 2011). It is an absolute quantification method which, unlike qPCR, does not rely on standard curves to estimate target DNA concentration. For DNA samples extracted from environmental water and containing potentially very low copy number of target DNA, the ability of ddPCR to detect rare events in a reaction is of particular interest, especially for invasive species in aquatic ecosystems where they may exist in low abundance and may be difficult to detect using conventional survey methods (e.g. netting and electrofishing).

Whether analysis is being carried out using qPCR or ddPCR, eDNA-based detection is becoming increasingly used in aquatic freshwater environments for a range of low abundance or invasive taxa, such as amphibians (Pilliod *et al.*, 2013, Spear *et al.*, 2014), molluscs (Goldberg *et al.*, 2013, Peñarrubia *et al.*, 2016, Carlsson *et al.*, 2017) and crustaceans (Tréguier *et al.*, 2014, Harper *et al.*, 2018), as well as fish (Takahara *et al.*, 2013, Klymus *et al.*, 2015, Davison *et al.*, 2016) – including salmonid species (Gustavson *et al.*, 2015, Atkinson *et al.*, 2018, Rusch *et al.*, 2018). To date, there has been no published studies for implementing eDNA for the detection of *O. gorbuscha.* We hypothesise that eDNA can be used for detection of non-native *O. gorbuscha* in running water. In addition, should any eradication measures be employed in the future, eDNA methods could be used to determine the efficacy of such efforts (Banks *et al.*, 2015).

This study aimed to address questions relating to the presence and distribution of *O. gorbuscha* in a Norwegian river, as part of a pilot study employing the non-intrusive and genetic survey method of eDNA collection and analysis. A species-specific probe-based assay was developed for this species and deployed in an urban river system where *O. gorbuscha* had been previously observed. We also developed this assay with the intention that it can be deployed in other European freshwater or marine ecosystems where this species may currently, or potentially, invade. This is of particular relevance for detection of the species during spawning season of this year (Autumn 2019), where it is unknown if the putative increased mortality of *O. gorbuscha* at sea (resulting from a mismatch between emigration time and food availability; Armstrong *et al.*, 2018) will result in a less significant invasion.

## MATERIALS AND METHODS

### Water sampling and eDNA extraction

For this study, water samples were collected from the Lysakerelva river in the south-east region of Norway (Table 1 and Figure 2) during Autumn of 2017 and Spring/early Summer of 2018. This river system runs through a large urban area outside of Oslo and is characterised by a number of natural migration barriers, including waterfalls. The smallest of these waterfalls (Møllefoss; Figure 2) is a few metres in height and is fitted with fish ladders to enable upstream migration. The next waterfall, Granfoss (Figure 2), is sufficiently high (∼12-15 metres) as to completely prevent upstream fish migration. In terms of the diversity of fish found in the study area, the Lysakerelva river fish fauna is dominated by *S. salar, S. trutta* and the European minnow *Phoxinus phoxinus* (Saltveit *et al.*, 2013). Several *O. gorbuscha* have previously been caught in the Lysakerelva river (>20 fish; Sandlund *et al.*, 2018).

**Table 1:**
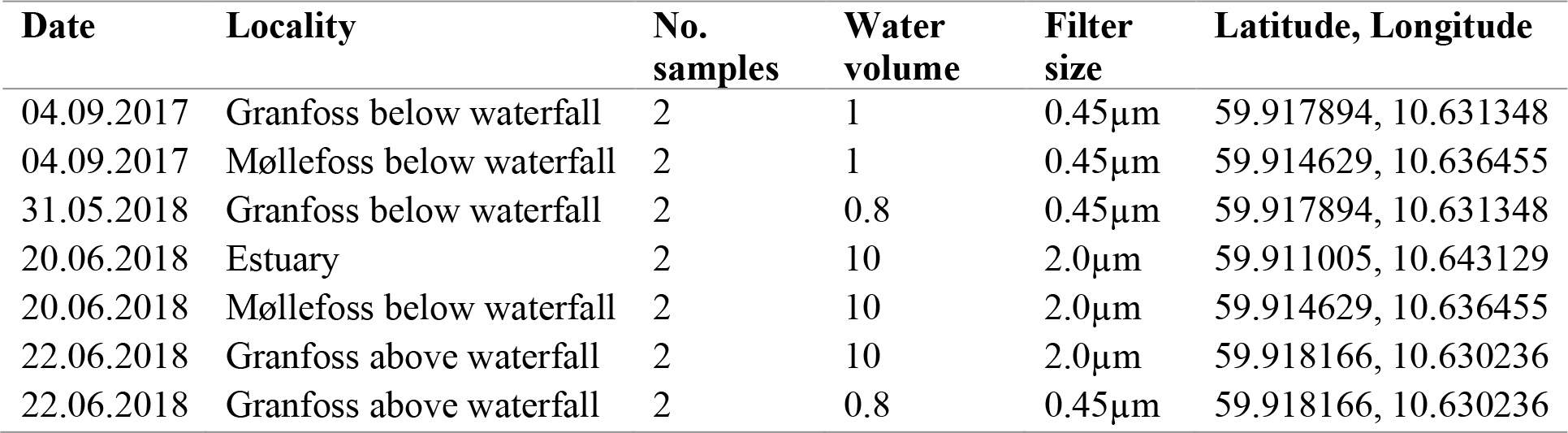
Details of water sampling in Lysakerelva river.

**Figure 2:**
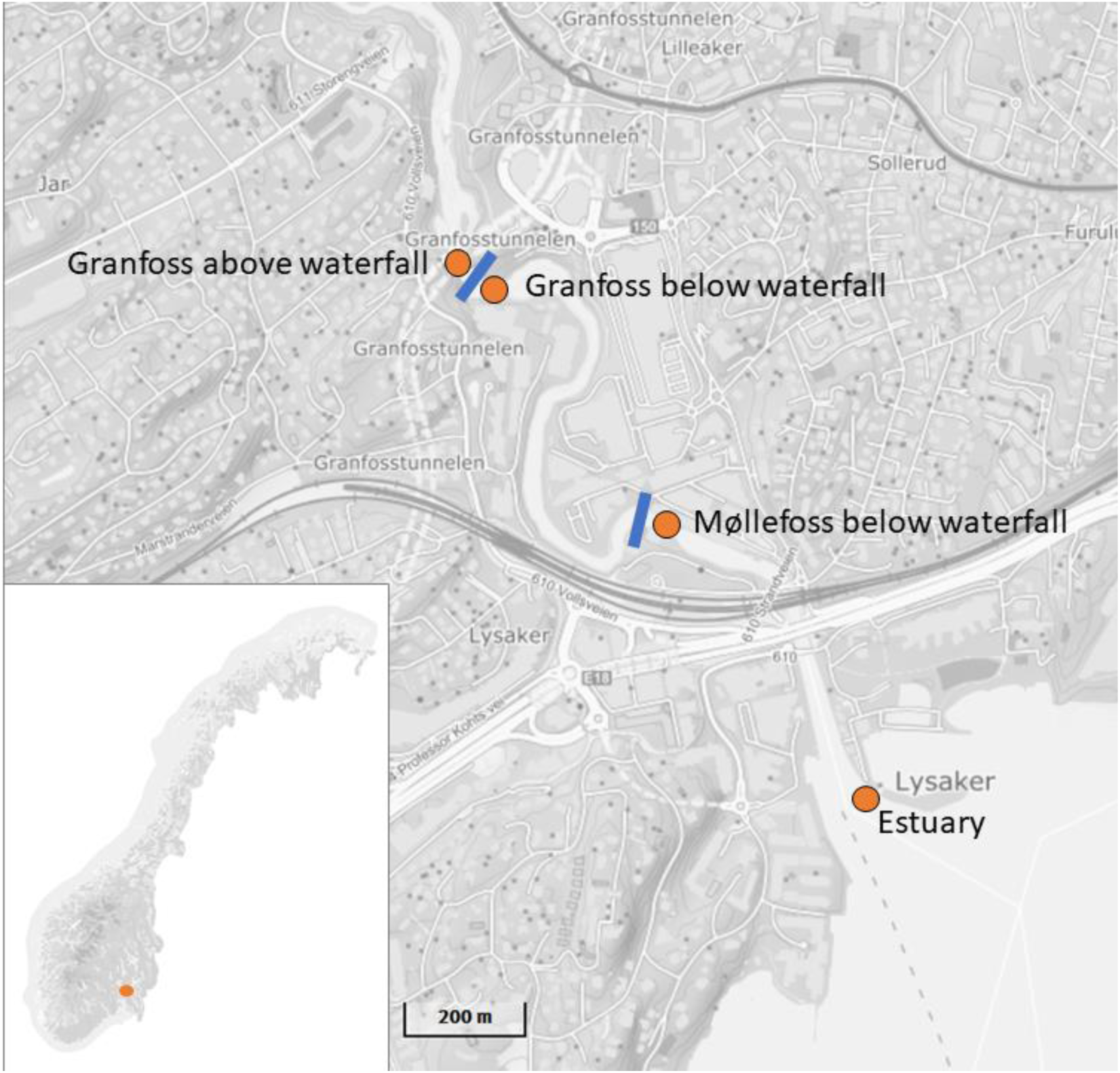
Boxplot showing the number of positive droplets from ddPCR detection of *O. gorbuscha* in relation to date and locality. The horizontal dashed line indicates the lower threshold of three droplets for assessing a sample as positive. See Table 1 for details of each sample.

During September 2017, two water samples were taken below Møllefoss and just above upper high tide close to the mouth of the Lysakerelva river, and two water samples were taken immediately below the Granfoss waterfall (∼800 metres upstream from the river mouth; Table 1 and Figure 2). These samples were taken to coincide with the spawning season of *O. gorbuscha.* This sampling was repeated the following May, the time period when it is expected that juvenile *O. gorbuscha* would be undertaking seaward migration (Table 1). In June 2018, two water samples were also taken from the estuary (Figure 2) as it was considered possible that *O. gorbuscha* may also be found in the coastal area after migrating downriver (Heard 1991; Moore *et al.*, 2016). Additional samples (*n*=4; Table 1) were taken from above the Granfoss waterfall in June 2018 (Figure 2) as negative eDNA field control samples, as it was expected that no *O. gorbuscha* would be present at this location due to the barrier of the Granfoss waterfall.

For each locality, we filtered two replicate samples of either *c*. 1 L water on a 0.45 μm cellulose filter (Pall MicroFunnel 300 ST; Pall Corporation, New York, USA; Table 1) or two replicate samples of *c*. 10 L water on a 2.0 μm glassfiber filter (Merck Millipore, Burlington, Massachusetts, USA; Table 1) using a peristaltic pump (Vampire sampler, Bürkle, Bad Bellingen, Germany). The 0.45 μm cellulose filters were immediately placed in 2 mL tubes with 1440 μL ATL-buffer (Qiagen, Hilden, Germany), whereas the 2.0 μm glassfiber filters were placed in 5 mL tubes with 4050 μL ATL-buffer. All samples were stored at room temperature until further processing in the genetics laboratory. All field equipment (e.g. filtering tubes and collection bottles) was sterilised between collection of each sample using 10% bleach solution for approximately 60 minutes.

In the laboratory, 160 μL or 450 μL Proteinase-K (Qiagen) was added to the 2 mL and 5 mL sampling tubes, respectively. All samples were incubated overnight at 56°C. DNA was isolated from 0.45 μm cellulose filters using DNeasy DNA Blood & Tissue kit (Qiagen), and from the 2.0 μm glassfiber filters using NucleoSpin Plant II Midi kit (Macherey-Nagel, Düren, Germany), following the manufacturers protocol except that Qiagen buffers were used instead of those supplied with the kit. DNA extracted from the 0.45 μm cellulose filters was eluted in 100 μL AE buffer, whereas DNA extracted from the 2.0 μm glassfiber filters was eluted in 200 μL AE buffer. All samples were re-eluted in order to maximise the output of DNA. Final concentrations of the eluted DNA samples (Supplementary Table 1) were determined using a Nanodrop 1000 Spectrophotometer (Thermo Fisher Scientific, Waltham, Massachusetts, USA).

### Molecular assay development and specificity testing

An assay was designed to amplify a 98 bp region of the mitochondrial *COI* gene of *O. gorbuscha* (Table 2). This assay consisted of primers and a 5’ VIC labelled TaqMan® minor groove binding (MGB) probe. The primers and probes were designed using Primer3 software (Rozen and Skaletsky 2000). Specificity of the assay was checked *in-silico*, by aligning the primers and probe with the consensus sequence generated from publicly available *O. gorbuscha* sequences as well as those from other salmonids commonly occurring in the study area (e.g. *S. salar* and *S. trutta*). The primer and probe sequences were also checked against the NCBI database to ensure specificity to the target organism. Furthermore, specificity of the assay was checked using qPCR, with tissue-extracted DNA from *O. gorbuscha*, as well as from *S. salar* and *S. trutta*. Tissue-extracted DNA was also acquired and tested by qPCR for rainbow trout (*O. mykiss*) which, while not a native salmonid, has been widely introduced throughout Europe (Stankovic *et al.*, 2015). In addition, DNA extracted from the closely-related chum salmon *O. keta* tested to check the specificity of the assay. The latter is not currently found in Europe but this species overlaps with *O. gorbuscha* in its native range.

**Table 2:**
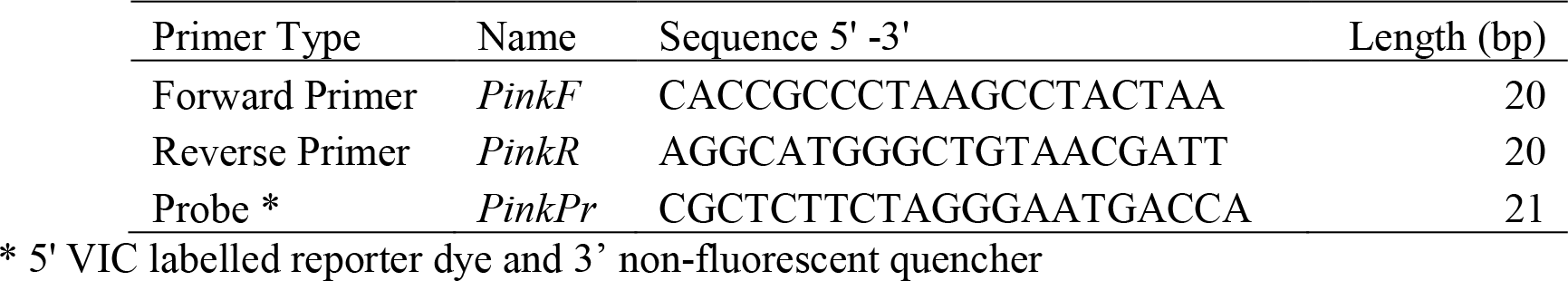
Details of the assay that was designed, tested and deployed for detection of a 98bp region of the mitochondrial *COI* gene of *O. gorbuscha* in this study.

All qPCR reactions took place in a 20 μl reaction volume, containing 10 μl of TaqMan™ Environmental Master Mix 2.0 (Thermofisher), 2 μl of each primer (2 μM), 2 μl of probe (2 μM; Applied Biosystems) and 2 μl of template DNA (where extracts were normalised to 34 ng/μl). The PCR program consisted of 50°C for 2 min, 95°C for 10 min, followed by 40 cycles of 95°C for 15 s and 60°C for 1 min. All qPCR reactions were carried out using QuantStudio™ 7 Flex Real-Time PCR System (Applied Biosystems). All qPCR analysis took place in University College Dublin.

Potential cross-amplification of other salmonids (*O. mykiss, S. salar, S. trutta, Salvelinus alpinus, Salvelinus fontinalis, Salvelinus namaycush and Thymallus thymallus*) using the *O. gorbuscha* assay was also tested using ddPCR at NINA using the same PCR conditions as those detailed in the next section.

### Digital droplet (dd)PCR analysis of eDNA samples

Sample collection and eDNA extraction resulted in a total of 14 samples originating from the Lysakerelva river (Table 1). Detection and concentration of target-DNA was assessed using droplet-digital-PCR (QX200 Droplet Digital PCR system with AutoDG, Bio-Rad Laboratories, Hercules, USA). A tissue-extracted DNA sample of *O. gorbuscha* was included in the analysis, as a positive control. A no-template control was also included in the analysis. The eDNA samples, along with positive, negative and no-template controls, were analysed in triplicate with the exception of the samples from 2017 (Table 1), which were analysedsingularly. The ddPCR reactions consisted of 0.9 μM forward and reverse primers, 0.25 μM of the probe, ddPCR™ Supermix for Probes (No dUTP) (Bio-Rad Laboratories), dH2O and 5 μL template. In order to assess potential problems with inhibition in our samples we also reran the samples with only 1 μL template. However, we found no indications of inhibition among our samples. To generate droplets, an AutoDG Instrument (Bio-Rad Laboratories) was used, with subsequent PCR amplification in a Veriti96-Well Thermal Cycler (Applied Biosystems). The following thermal cycling conditions were used: an initial denaturation step at 95°C for 10 min, 40 cycles of denaturation at 95°C for 30 sec, annealing and extension at 60°C for 1 min, a final step of denaturation at 98°C for 10 min, and a final hold at 4°C. PCR plates were transferred to a QX200 Droplet Reader (Bio-Rad Laboratories) to automatically detect the fluorescent signal in the droplets. QuantaSoft software v.1.7.4 (Bio-Rad Laboratories) was used to separate positive from negative droplets according to manufacturer’s instructions. A threshold of minimum 3 positive droplets were used as a criterion for positive detection Wacker et al. 2019). All ddPCR-analyses took place at the genetic lab at NINA in Trondheim, Norway.

## RESULTS

### Molecular assay development

Testing of the *O. gorbuscha* assay with non-target salmonids using tissue-extracted DNA and qPCR, revealed that the assay did not cross-amplify *S. salar, S. trutta*, or *O. mykiss.* ddPCR testing of non-target salmonid species (*O. mykiss, S. salar, S. trutta, S. alpinus, S. fontinalis, S. namaycush, T thymallus*) did not produce any detectable amplification. However, amplification was observed for *O. keta* DNA analysed by qPCR (data not shown). This species is closely related to *O. gorbuscha* and there are very few nucleotide differences between these species for the *COI* region targeted by our assay (Supplementary Figure 3). However, this should not present a major concern for researchers wishing to apply this assay for *O. gorbusha* detection in Norwegian and other European locations, as neither *O. keta* nor other species of Pacific salmon (*O. nerka, O. kisutch, O. tshawytscha* and *O. masoudo*) currently occur in these regions.

### ddPCR analysis of eDNA samples

An average number of 15,299 droplets were analysed for each sample included in the ddPCR analyses. Using the *O. gorbuscha* assay in a ddPCR analysis, there was target DNA detected in six out of 14 eDNA samples (Table 3).

**Table 3:** Detection of *O. gorbuscha* using eDNA shown as the number of samples that were positive for detection out of the total number of samples analysed by ddPCR.

| Locality | Date | eDNA detection |
| --- | --- | --- |
| Granfoss below waterfall | 04.09.2017 | 1/2 |
| Granfoss below waterfall | 31.05.2018 | 2/2 |
| Granfoss above waterfall | 22.06.2018 | 1/4 |
| Møllefoss below waterfall | 04.09.2017 | 2/2 |
| Møllefoss below waterfall | 20.06.2018 | 0/2 |
| Estuary | 20.06.2018 | 0/2 |

In Autumn 2017, *O. gorbuscha* DNA was detected at Granfoss but not at Møllefoss (Table 3, Figure 3). In Spring, target DNA was detected in both samples taken below the Granfoss waterfall, which indicates that there were still young *O. gorbuscha* in the sampling area that had not yet left the river. No target DNA was found in samples taken near to the mouth of the river in June 2018, which is perhaps unsurprising, considering that upstream and downstream samples were taken 20 days apart and young *O. gorbuscha* may have already migrated to sea.

**Figure 3:**
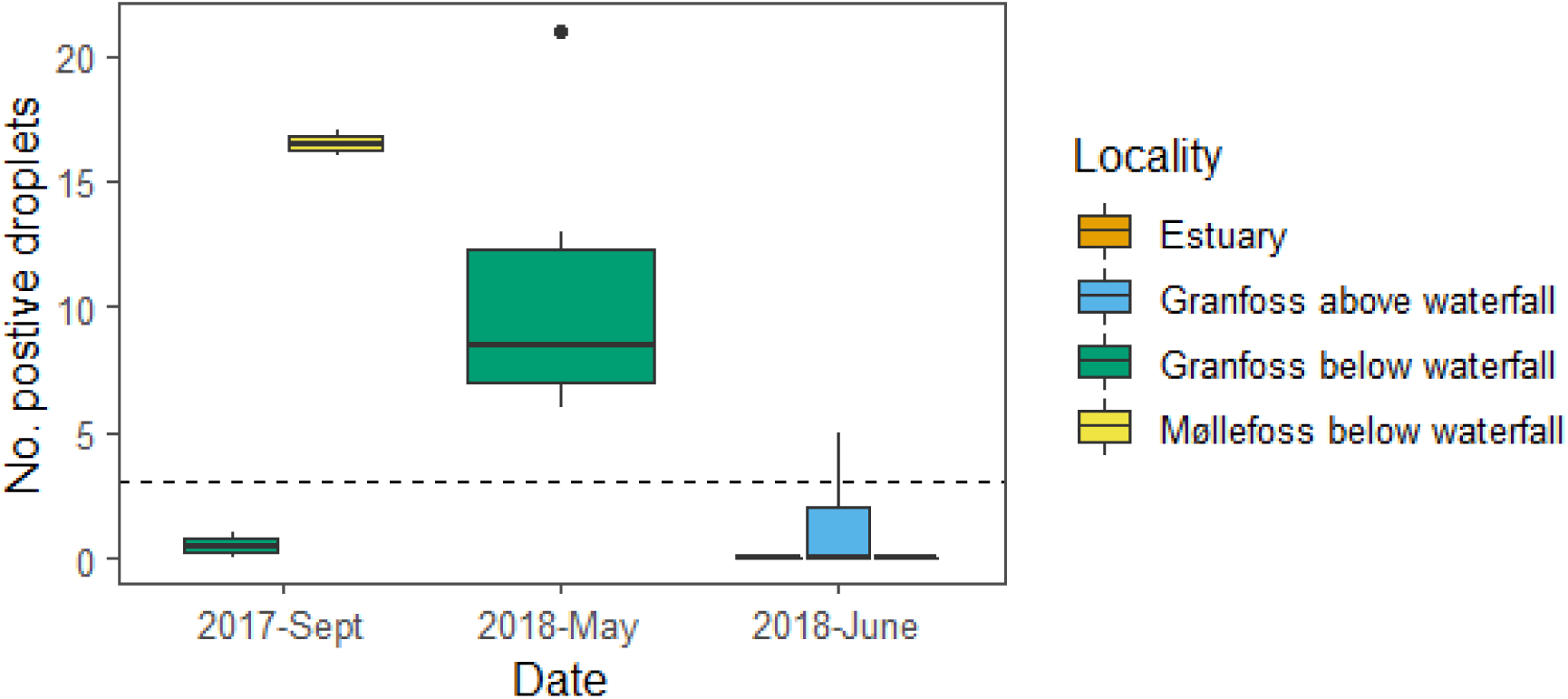
Boxplot showing the number of positive droplets from ddPCR detection of *O. gorbuscha* in relation to date and locality. The horizontal dashed line indicates the lower threshold of three droplets for assessing a sample as positive. See Table 1 for details of each sample.

No detectable target DNA was found in samples taken from the estuary. It is possible that, similar to the samples taken in the downstream region of the Lysakerelva river, young *O. gorbuscha* may have already migrated out to sea when this sampling took place in the Summer. Alternatively, a lack of detection of *O. gorbuscha* at this location may be a false negative, as a result of the relatively low number of samples taken over a large sampling area (Table 1 and Figure 2).

Interestingly, target DNA was also detected in low copy number from one out of four of the control samples above the migration barrier (Figure 3) indicating alternative means of target DNA transport to this site, or contamination of the sample in the field (see Discussion). All no-template controls were negative for target DNA.

## DISCUSSION

The year 2017 was unprecedented in terms of the number of *O. gorbuscha* that were observed in Norwegian rivers (Mo *et al.*, 2018), as well as in Ireland (Whelan 2017, Millane *et al.*, 2019) and Scotland (Armstrong *et al.*, 2018,). In the present study, we were successfully able to detect *O. gorbuscha* from environmental water samples in Norway during this invasion. Target DNA was amplified from samples taken in both Autumn (during adult spawning) and the following Spring (during the migration of juveniles), indicating that spawning in the focal river system was successful. However, the survival of juvenile *O. gorbuscha* at sea is poorly understood and it is unknown whether mortality will limit the ability of this species to return to the river and establish self-reproducing populations.

The assay developed and validated in this study not only has applications for monitoring presence of *O. gorbuscha* spatially and temporally in Norwegian rivers, but also in countries where it currently exists outside of its native range, as well as those where it has not been observed but may potentially occur in the future. Further, this assay can be used to monitor *O. gorbuscha* presence following any future eradication efforts that may be implemented. While the utility of the assay developed in this study is limited to those areas where *O. gorbuscha* does not overlap with *O. keta*, this should not be of concern for researchers using this assay to detect *O. gorbuscha* within its invasive range as *O. keta* is not currently found outside the North Pacific.

The finding of a positive detection of *O. gorbuscha* above the impassable barrier of Granfoss waterfall demonstrates the potential drawbacks of using eDNA to infer species presence. There was no indication of contamination in any of the negative controls included in this study. Specificity testing by qPCR and ddPCR revealed that this assay does not amplify *COI* from other commonly co-occurring native or introduced salmonids. The sample location was the only site visited on the day of sampling, yet it is impossible to rule out contamination of field equipment from previous sampling events downstream of the barrier. It is also possible that an alternative vector, such as predation (by birds for example; Merkes *et al.*, 2014), can explain the detection of target DNA in a location where the organism is not present.

The results of our pilot study show that this assay can be used to detect the presence of *O. gorbuscha* in running water. However, we can make some suggestions to increase the efficacy of our approach for future studies. The lack of detection of *O. gorbuscha* in water samples taken from the estuary and at the mouth of the Lysakerelva river in early Summer indicated that the juveniles had already migrated to sea. We would recommend an increase in temporal sampling, as data derived from these samples may have been able to reveal the timing of juvenile migration to sea, increasing our knowledge about *O. gorbuscha* behaviour in Norwegian rivers and indicating the extent of potential interactions with local fauna.

We detected a mismatch in the forward primer sequence (Supplementary Figure 1), based on the *O. gorbuscha* consensus sequence generated from publicly available *COI* records on GenBank. The source of this mismatch is a sequence from an odd-year individual (Accession No. MG951587.1) from the White Sea that was submitted to the NCBI database after the assay was developed and tested. It is therefore possible that this haplotype is found in Norwegian rivers. This mismatch is found on the 5’ end of the forward primer, therefore it is unlikely that our assay efficiency was severely compromised in the present study. However, to ensure optimal efficiency in the assay (and particularly to ensure sensitive detection of the low copy numbers frequently found in eDNA samples), we recommend that the forward primer incorporate a degenerate base at this position should the assay be deployed in other studies. Further, we concur with other eDNA researchers (e.g. Goldberg *et al.*, 2016) that concerted efforts are made to minimise any contamination in the field, through the implementation of careful sterilisation and sampling techniques, as well as the use of strict controls (e.g. field blanks, filter blanks) to monitor for exogenous sources of DNA during the entire eDNA sampling and analysis workflow.

## ACKNOWLEDGMENTS

The authors would like to thank Hege Brandsegg for running the ddPCR genetic analysis. We also gratefully acknowledge Sigrid Skoglund for drawing the pink salmon life cycle.

## COMPLIANCE WITH ETHICAL STANDARDS

### Ethical approval

All applicable international, national, and/or institutional guidelines for the care and use of animals were followed.

