## Supplementary Table 1 for "Environmental (e)DNA detection of the invasive pink salmon *Oncorhynchus gorbuscha* during the 2017 Norwegian invasion"

**Supplementary Table 1:** Concentrations with 260/280 and 260/230 ratios resulting from spectrophotometer analysis of environmental DNA samples. For sample details see main text Table 1.

| **Sample no.** | **Filter size** | **DNA isolation kit** | **Elution volume** | **OD** | **260/280** | **260/230** |
|---|---|---|---|---|---|---|
| **1** | 0.45µm | Qiagen Blood & Tissue | 100 | 46.99 | 1.74 | 0.78 |
| **2** | 0.45µm | Qiagen Blood & Tissue | 100 | 49.31 | 1.73 | 0.76 |
| **3** | 0.45µm | Qiagen Blood & Tissue | 100 | 40.72 | 1.81 | 0.84 |
| **4** | 0.45µm | Qiagen Blood & Tissue | 100 | 50.93 | 1.73 | 0.76 |
| **5** | 2.0µm | Machery-Nagel NucleoSpin Plant II Midi | 200 | 305.78 | 1.78 | 1.47 |
| **6** | 2.0µm | Machery-Nagel NucleoSpin Plant II Midi | 200 | 280.49 | 1.76 | 1.44 |
| **7** | 2.0µm | Machery-Nagel NucleoSpin Plant II Midi | 200 | 61.56 | 1.75 | 1.35 |
| **8** | 2.0µm | Machery-Nagel NucleoSpin Plant II Midi | 200 | 57.37 | 1.7 | 1.36 |
| **9** | 0.45µm | Qiagen Blood & Tissue | 100 | 29.18 | 1.86 | 1.69 |
| **10** | 0.45µm | Qiagen Blood & Tissue | 100 | 45.49 | 1.86 | 1.57 |
| **11** | 0.45µm | Qiagen Blood & Tissue | 100 | 64.09 | 1.82 | 1.58 |
| **12** | 1.2µm | Qiagen Blood & Tissue | 100 | 64.67 | 1.75 | 1.34 |
| **13** | 2.0µm | Machery-Nagel NucleoSpin Plant II Midi | 200 | 73.65 | 1.67 | 1.35 |
| **14** | 2.0µm | Machery-Nagel NucleoSpin Plant II Midi | 200 | 74.11 | 1.70 | 1.47 |
