## Supplementary Figure 1 for "Environmental (e)DNA detection of the invasive pink salmon *Oncorhynchus gorbuscha* during the 2017 Norwegian invasion"

```
PinkF         1 CACCGCCCTAAGCCTACTAA------------------------------------------------------------------------------ 98
PinkR         1 ------------------------------------------------------------------------------AATCGTTACAGCCCATGCCT 98
PinkPr        1 ---------------------------------------------CGCTCTTCTAGGGAATGACCA-------------------------------- 98
O. gorbuscha  1 CACCGCCMTAAGCCTACTAATTCGGGCAGAACTAAGCCAGCCAGGCGCTCTTCTAGGGAATGACCAGATCTATAACGTAATCGTTACAGCCCATGCCT 98
O. mykiss     1 YACCGCCCTGAGTCTAMTGATTCGRGCGGAMYTAAGCCWSCMGGGMKYYYTTCTRRSGGATRACYMWAWCTATAACGTGATCGTCACAGCCCATGCCT 98
O. keta       1 CACCGCCCTRAGCCTACTRATTCGGGCAGAACTAAGCCAGCCAGGCGCTCTTCTRGGRGATGACCAGATCTAYAAYGTAATCGTYACAGCCCATGCCT 98
S. alpinus    1 CACCGCCCTTAGCCTYTTGATCCGGGCAGAGTTAAGCCAACCCGGAGCTCTTCTAGGGGATGACCAGATCTATAACGTAATCGTAACAGCCCATGCCT 98
S. trutta     1 CACCRCCYTAAGTCTCTTRATTCGGGCMGAACTYAGCCAACCCGGCGCYCTCCTAGGGGATGACCAGATYTATAACGTAATYGTWACAGCCCATGCCT 98
S. salar      1 CACCGCCCTAAGTCTCTTGATTCGAGCAGAACTCAGCCAGCCTGGCGCCCTTCTGGGAGATGACCAAATTTATAACGTAATTGTTACRGCCCATGCCT 98
```

**Supplementary Figure 1:** Primers (*PinkF* and *PinkR*) and probe (*PinkPr*) designed for targeted detection of *O. gorbuscha* are shown aligned to the target consensus sequence, as well as consensus sequences for other salmonid species. Mismatches are highlighted by shading.
